# Oxygen supply limits the heat tolerance of locusts during the first instar only

**DOI:** 10.1101/2020.01.16.909705

**Authors:** Jacob P. Youngblood, John M. VandenBrooks, Oluwatosin Babarinde, Megan E. Donnay, Deanna B. Elliott, Jacob Fredette-Roman, Michael J. Angilletta

## Abstract

Extreme heat directly limits an organism’s survival and reproduction, but scientists cannot agree on what causes organisms to lose function or die during heating. According to the theory of oxygen- and capacity-limitation of thermal tolerance, heat stress occurs when a warming organism’s demand for oxygen exceeds its supply, triggering a widespread drop in ATP concentration. This model predicts that an organism’s heat tolerance should decrease under hypoxia, yet most terrestrial organisms tolerate the same amount of warming across a wide range of oxygen concentrations. This point is especially true for adult insects, who deliver oxygen through highly efficient respiratory systems. However, oxygen limitation at high temperatures may be more common during immature life stages, which have less developed respiratory systems. To test this hypothesis, we measured the effects of heat and hypoxia on the survival of locusts (*Schistocerca cancellata*) throughout development. We demonstrate that the heat tolerance of locusts depends on oxygen supply during the first instar but not during later instars. This finding provides further support for the idea that oxygen limitation of thermal tolerance depends on respiratory performance, especially during immature life stages.

## 1. Introduction

Pörtner and colleagues argued that heat tolerance depends on an organism’s ability to deliver oxygen during warming (Pörtner 2002; Pörtner et al. 2010), but this idea has been controversial among comparative physiologists (Grans et al., 2014; Jutfelt et al., 2018; Pörtner and Giomi, 2013; Pörtner et al., 2017). Recent reviews emphasized that oxygen delivery is just one of many processes that can fail during heat stress; the particular process that fails for an organism will depend on its life history, activity level, and respiratory performance (Clark et al., 2013; Gangloff and Telemeco, 2018; Macmillan, 2019; Schulte, 2015). For example, oxygen supply limits the heat tolerance of aquatic insects more severely than that of terrestrial insects (reviewed by Verberk et al. 2016). This difference arises partly because terrestrial insects live in air, where oxygen is abundant and easy to transport (Giomi et al., 2014; Verberk et al., 2013). However, even air-breathing insects can experience oxygen limitation during immature life stages where oxygen supply depends on a respiratory system that has yet to develop fully (Harrison et al., 2017).

Embryos are arguably the most vulnerable to oxygen limitation, because their oxygen delivery depends on diffusion across the eggshell (Woods, 2010). As temperature rises, an embryo’s metabolic demand can increase more quickly than its rate of diffusion (Woods and Hill, 2004). After hatching, larvae still rely on diffusion to deliver oxygen, but it now occurs through the tracheal system, a series of hollow tubes connecting the tissues and atmosphere through spiracles. Diffusion alone can supply the oxygen needed to meet the metabolic demands of small larvae, but larger larvae and adults regulate pressure gradients within the tracheal system to move oxygen by convection (Weis-Fogh, 1964a; Weis-Fogh, 1964b; Weis-Fogh, 1967). For instance, grasshoppers (*Schistocerca americana)* generate convective airflow with abdominal muscle contractions that repeatedly compress and expand the tracheae (Harrison et al. 2013). Grasshoppers use abdominal pumping to enhance oxygen delivery during metabolic challenges such as heat, hypoxia, or activity (Harrison et al., 2006; Lighton and Lovegrove, 1990; Snelling et al., 2017), but this ventilatory response does not develop until the third instar (Greenlee and Harrison, 2004a; Lee et al., 2013). Therefore, oxygen delivery becomes easier as grasshoppers advance from instar-to-instar. (Greenlee and Harrison, 2004b; Greenlee et al., 2009; Harrison et al., 2005; Hartung et al., 2004; Kirkton et al., 2005; Lease et al., 2006). If the heat tolerance of a grasshopper depends on its capacity to deliver oxygen, these ontogenetic changes in ventilation should make them more susceptible to heat and hypoxia during early life stages.

Previously, researchers found that some life stages are more susceptible to heat and hypoxia than others. For example, larval beetles (*Tenebrio molitor*) died from heat stress 10% faster in hypoxia than normoxia, but adult beetles died at the same rate in either oxygen concentration (Mccue et al., 2015). Similarly, hypoxia reduced the heat tolerance of silk moth larvae (*Bombyx mori*) but not that of pupae (Boardman et al., 2015). Verberk and colleagues (2013) proposed that such discrepancies stem from different ways that organisms acquire oxygen; organisms that can upregulate oxygen delivery as supply increases should tolerate heat and hypoxia better than organisms that cannot. To test this hypothesis, we measured the effects of heat and hypoxia on the survival and development of South American locusts (*Schistocerca cancellata*) from hatching to adulthood and from the beginning to end of the 1^st^, 3^rd^, and 5^th^ instars. We predicted that heat and hypoxia would interact to kill more locusts than heat and hypoxia alone, especially during the 1^st^ instar when oxygen delivery depends primarily on diffusion.

## 2. Materials and methods

Locusts were obtained from a breeding colony at Arizona State University, established from a sample of more than 1000 individuals, collected near Casa de Piedras of Argentina in 2015 (Medina et al., 2017). Locusts were housed in an aluminium-framed screen cage (45 × 45 × 45 cm) within an environmental chamber (Conviron, Winnipeg, Canada) programmed to control light and temperature. The light cycle was 14L:10D, and the temperatures were 35 °C during the day and 25 °C at night. A 75-watt infrared lamp generated heat for behavioural thermoregulation. Locusts were fed an excess of lettuce, wheat bran, and wheat grass daily. During the experiments, locusts were housed individually in plastic cups (946 mL) with a perch to facilitate molting. We supplied fresh food daily and removed uneaten food and frass every other day.

For the first experiment, we collected 96 locusts from the colony on the first day of their first instar. We reared these locusts from hatching to adulthood in one of four treatments: cold and normoxic (a diel cycle of 21°-35°C and 21% O_2_); hot and normoxic (a diel cycle of 28°-42°C and 21% O_2_); cold and hypoxic (a diel cycle of 21°-35°C and 13% O_2_); and hot and hypoxic (a diel cycle of 28°-42°C and 13% O_2_). To control atmospheric oxygen, we put the containers of locusts in each treatment into a chamber (60 × 45 × 37 cm; SKU# 1006267, The Container Store, USA). The atmosphere in each chamber was regulated by a ROXY-4 Universal Controller (Sable Systems International, Las Vegas, NV, USA). To control temperature, each chamber was placed in an incubator (DR-36VL; Percival Scientific, Perry, Iowa, USA), programmed to maintain the diel cycle of temperature. We recorded daily survival, the time to reach the 2nd instar, and the time to reach adulthood.

We repeated the experiment with locusts collected at the beginning of the third instar and locusts collected at the beginning of the fifth instar (N=65 and N=40, respectively). These locusts were followed until their next instar. We recorded daily survival and time to next instar. Locusts were scored as surviving to the next instar if they were alive 24 hours after molting.

We fit three statistical models to the data using the R statistical software (version 3.4.1; R Core Team 2017). First, we modeled the probability of locusts surviving specific instars in each treatment. To do so, we used the *nlme* library to fit a generalized linear model with a binomial distribution of error and fixed effects of temperature, oxygen, and instar. Second, we modeled the probability of locusts surviving each day, from hatching to adulthood. For this analysis, we used the *survival* library to fit a Cox proportional-hazards model with effects of temperature and oxygen. Finally, we modeled the development time of locusts during the first, third, and fifth instars. We used the *nlme* library to fit a generalized linear model with a Gaussian distribution of error and fixed effects of temperature, oxygen, and instar. For each of these analyses, we used the *MuMIn* library (Bartoń, 2013) to perform multimodel averaging. This package fits a set of models containing all possible subsets of the fixed factors and calculates the weighted average of each parameter value using the likelihood of the full model and all possible subsets of the full model. Model averaging reduces the bias in estimating sources of variation by using all likely models instead of one model with less than full likelihood (Burnham and Anderson, 1998; Harrison et al., 2018).

## 3. Results

A locust’s probability of surviving to the next instar depended on its body temperature, oxygen supply, and life stage. As shown in Table 1, the most likely models included main effects of temperature, oxygen, and instar, as well as some interactions between these variables. Based on means estimated by model averaging, third instar locusts would be more than 93% likely to survive in any treatment (Fig. 1). The probabilities of a locust surviving the third or fifth instar were similar among treatments. However, locusts in the first instar were more susceptible to death under heat and hypoxia. During this early life stage, only 67% of locusts survived to the next instar in the hot and hypoxic treatment, compared to 90% or greater survivorship in the other treatments.

**Table 1.** The most likely models of survival included effects of temperature, oxygen, and instar. Each model predicts the probability of surviving to the next instar. For each model, we report the number of parameters (*k*), the log likelihood, Akaike information criterion (AIC), and the Akaike weight (*w*), which is the probability that the model describes the data better than other models

| Model | $k$ | Log likelihood | AIC | ΔAIC | $w$ |
| --- | --- | --- | --- | --- | --- |
| temp + O<sub>2</sub> + instar + (temp · O<sub>2</sub>) + (temp · instar) | 8 | -56.7 | 130.2 | 0 | 0.26 |
| temp + O<sub>2</sub> + instar + (temp · O<sub>2</sub>) + (temp · instar) + (O<sub>2</sub> · instar) | 10 | -55.2 | 131.6 | 1.42 | 0.13 |
| temp + O<sub>2</sub> + instar + (temp · O<sub>2</sub>) + (O<sub>2</sub> · instar) | 8 | -57.5 | 131.9 | 1.67 | 0.11 |
| temp + O<sub>2</sub> + instar + (temp · O<sub>2</sub>) | 6 | -59.7 | 131.9 | 1.68 | 0.11 |
| temp + O<sub>2</sub> + instar + (temp · instar) + (O<sub>2</sub> · instar) | 9 | -56.9 | 133.0 | 2.76 | 0.06 |
| temp + O<sub>2</sub> + instar + (temp · instar) | 7 | -60.6 | 133.1 | 2.85 | 0.06 |
| O<sub>2</sub> + instar + (O<sub>2</sub> · instar) | 6 | -62.7 | 133.7 | 3.50 | 0.04 |
| O<sub>2</sub> + instar | 4 | -60.6 | 133.8 | 3.55 | 0.04 |
| temp + instar + (temp · instar) | 6 | -62.7 | 134.1 | 3.90 | 0.04 |
| temp + O<sub>2</sub> + instar | 5 | -62.1 | 134.6 | 4.36 | 0.03 |
| temp + O<sub>2</sub> + instar + (O<sub>2</sub> · instar) | 7 | -60.0 | 134.6 | 4.41 | 0.03 |
| instar | 3 | -64.3 | 134.8 | 4.64 | 0.03 |
| temp + O<sub>2</sub> + instar + (temp · O<sub>2</sub>) + (temp · instar) + (O<sub>2</sub> · instar) + (temp · O<sub>2</sub> · instar) | 12 | -54.5 | 134.9 | 4.71 | 0.02 |
| temp + instar | 4 | -63.7 | 135.7 | 5.45 | 0.02 |
| temp + O<sub>2</sub> + (temp · O<sub>2</sub>) | 4 | -64.1 | 136.5 | 6.31 | 0.01 |

**Figure 1.**
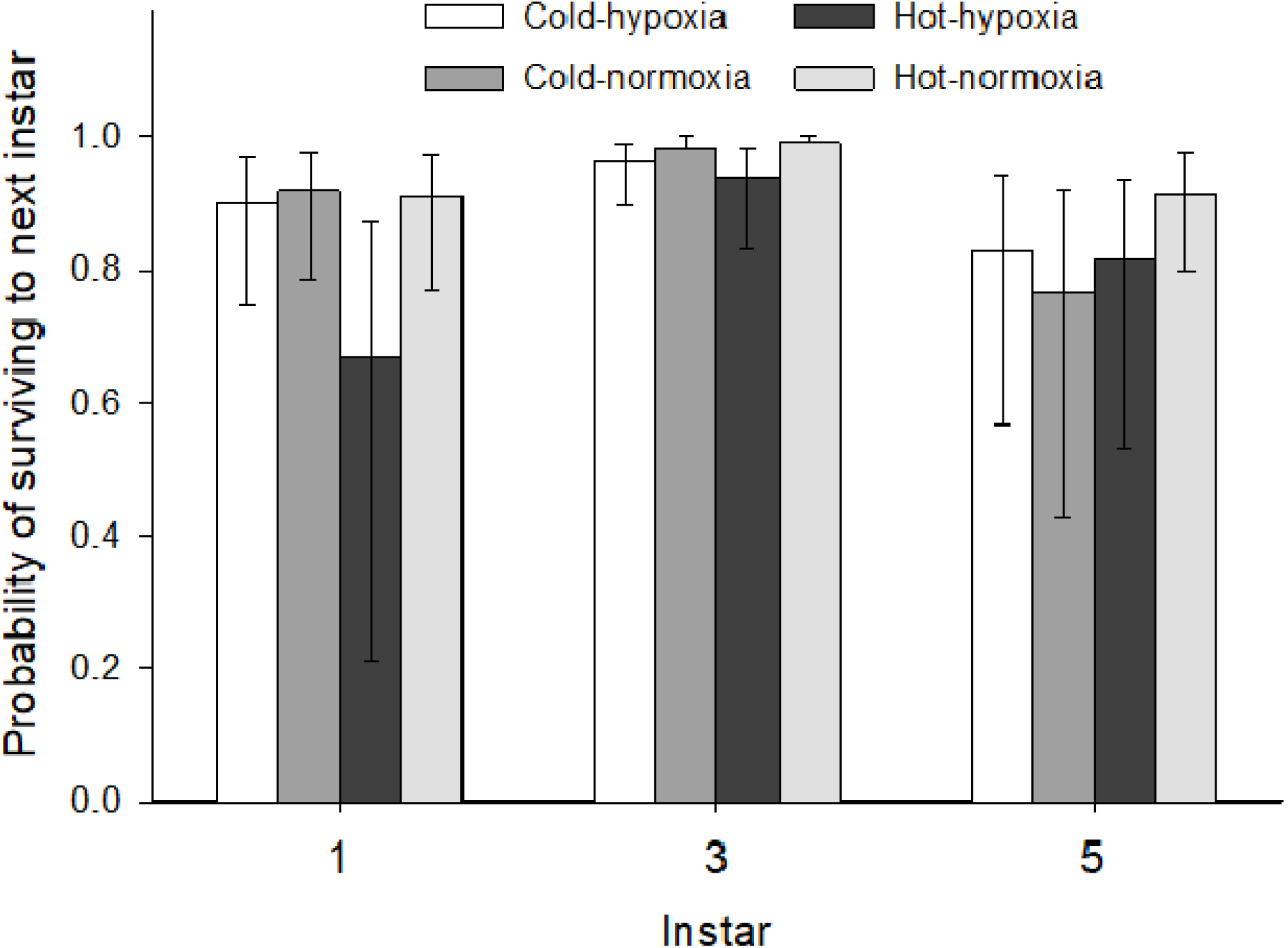
Heat and hypoxia interacted to kill first instar locusts. Bars denote the most likely probability of locusts surviving to the next instar, calculated from multimodel averaging, in cold or hot thermal regimes (21-35 or 28-42 °C, respectively), under either hypoxia or normoxia (13 or 21% oxygen, respectively). Error bars represent 95% confidence intervals.

Survivorship from hatching to adulthood depended on temperature more than oxygen concentration (Table 2). Temperature was included in all likely models, summing to a likelihood of 1 (Table S1). Specifically, locusts died more often in the treatments with higher temperatures (Fig. 2). Only 9 of 42 locusts in the hot treatments survived to adulthood, whereas 20 of 46 in the cold treatments survived to the same stage.

**Table 2.** The most likely model of survival during development included only an effect of temperature. Each model is a Cox proportional-hazard models that predict the probability of survival over time. For each model, we report the number of parameters (*k*), the log likelihood, Akaike information criterion (AIC), and the Akaike weight (*w*), which is the probability that the model describes the data better than other models

| Model | $k$ | Log likelihood | AIC | ΔAIC | $w$ |
| --- | --- | --- | --- | --- | --- |
| temperature | 2 | -204.8 | 411.8 | 0 | 0.56 |
| temperature + O<sub>2</sub> | 3 | -204.5 | 413.4 | 1.56 | 0.26 |
| temperature + O<sub>2</sub> + (temperature · O<sub>2</sub>) | 4 | -203.8 | 414.1 | 2.26 | 0.18 |
| O<sub>2</sub> | 2 | -220.8 | 443.7 | 31.9 | 0 |
| null | 2 | -220.9 | 445.8 | 34.0 | 0 |

**Figure 2.**
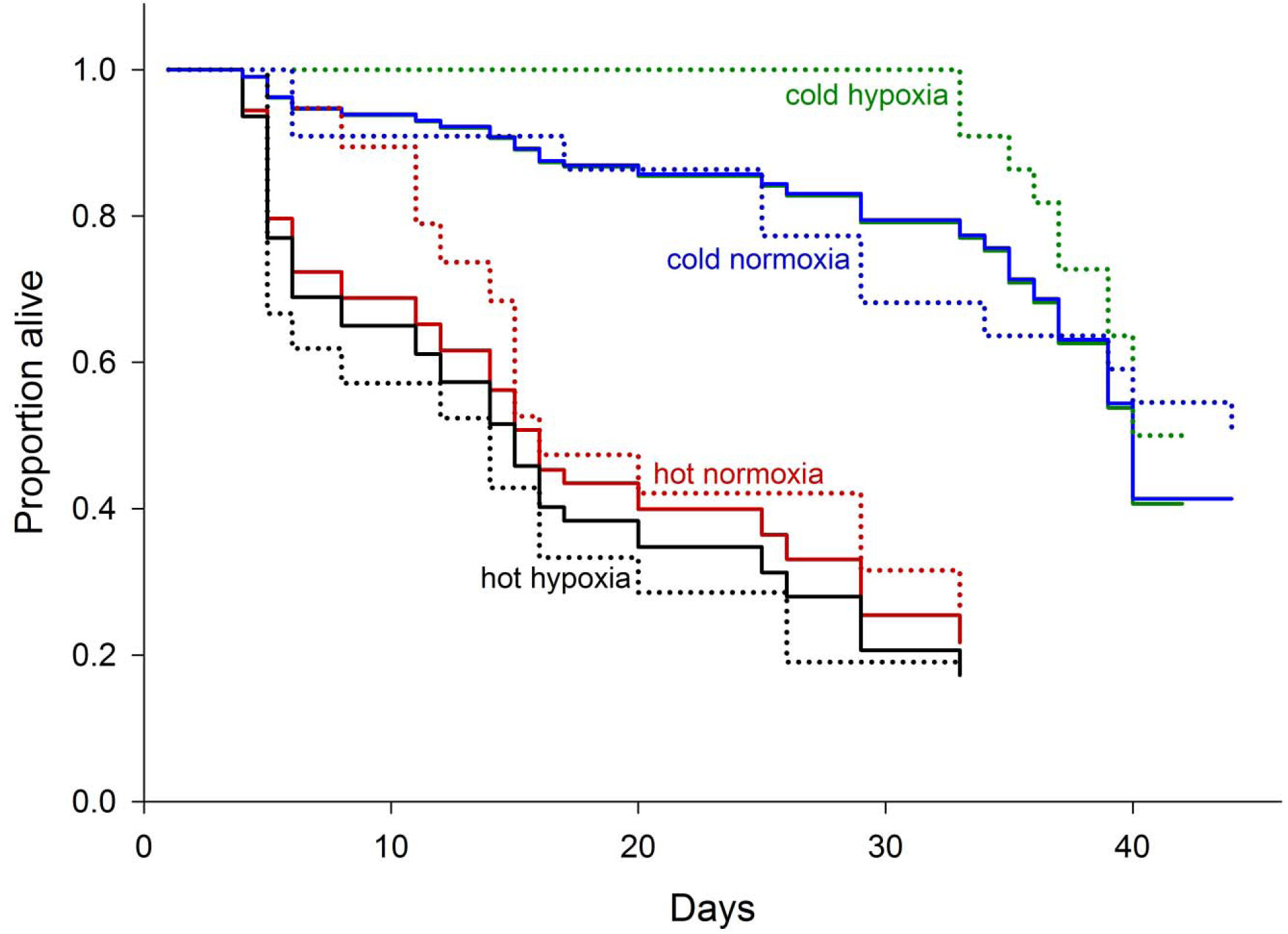
Locusts are more likely to die at high temperatures. Dotted lines denote the proportion of locusts that were alive each day of the experiment. Solid lines denote the most likely probability of a locust surviving each day, calculated from multi-model averaging, in cold or hot thermal regimes (21-35 or 28-42 °C, respectively) under either hypoxia or normoxia (13 or 21% oxygen, respectively).

Development time depended on temperature and instar more than oxygen concentration. (Table 3). Temperature, instar, and their interaction were included in all likely models, summing to a likelihood of 1. Oxygen concentration was less likely to be included in the best model of development time (Table S1). Not surprisingly, locusts of all instars developed faster in the warmer treatment. This effect was greatest during the fifth instar (Fig. 3). Across all treatments, locusts in the fifth instar (N = 38) took longer to develop than did locusts in either the first or third instar (N = 73 and 63, respectively).

**Table 3.** Temperature, oxygen, and instar affected development times. For each model, we report the number of parameters (*k*), the log likelihood, Akaike information criterion (AIC), and the Akaike weight (*w*), which is the probability that the model describes the data better than other models.

| Model | $k$ | Log likelihood | AIC | ΔAIC | $w$ |
| --- | --- | --- | --- | --- | --- |
| temp + O<sub>2</sub> + instar + (temp · O<sub>2</sub>) + (temp · instar) + (O<sub>2</sub> · instar) + (temp · O<sub>2</sub> · instar) | 13 | -209.2 | 446.6 | 0 | 0.36 |
| temp + O<sub>2</sub> + instar + (temp · O<sub>2</sub>) + (temp · instar) | 9 | -214.3 | 447.6 | 1.05 | 0.22 |
| temp + O<sub>2</sub> + instar + (temp · O<sub>2</sub>) + (temp · instar) + (O<sub>2</sub> · instar) | 11 | -212.3 | 448.2 | 1.62 | 0.17 |
| temp + instar + (temp · instar) | 7 | -216.9 | 448.5 | 1.95 | 0.14 |
| temp + O<sub>2</sub> + instar + (temp · instar) + (O<sub>2</sub> · instar) | 10 | -214.5 | 450.3 | 3.74 | 0.06 |
| temp + O<sub>2</sub> + instar + (temp · instar) | 8 | -216.8 | 450.5 | 3.92 | 0.05 |

**Figure 3.**
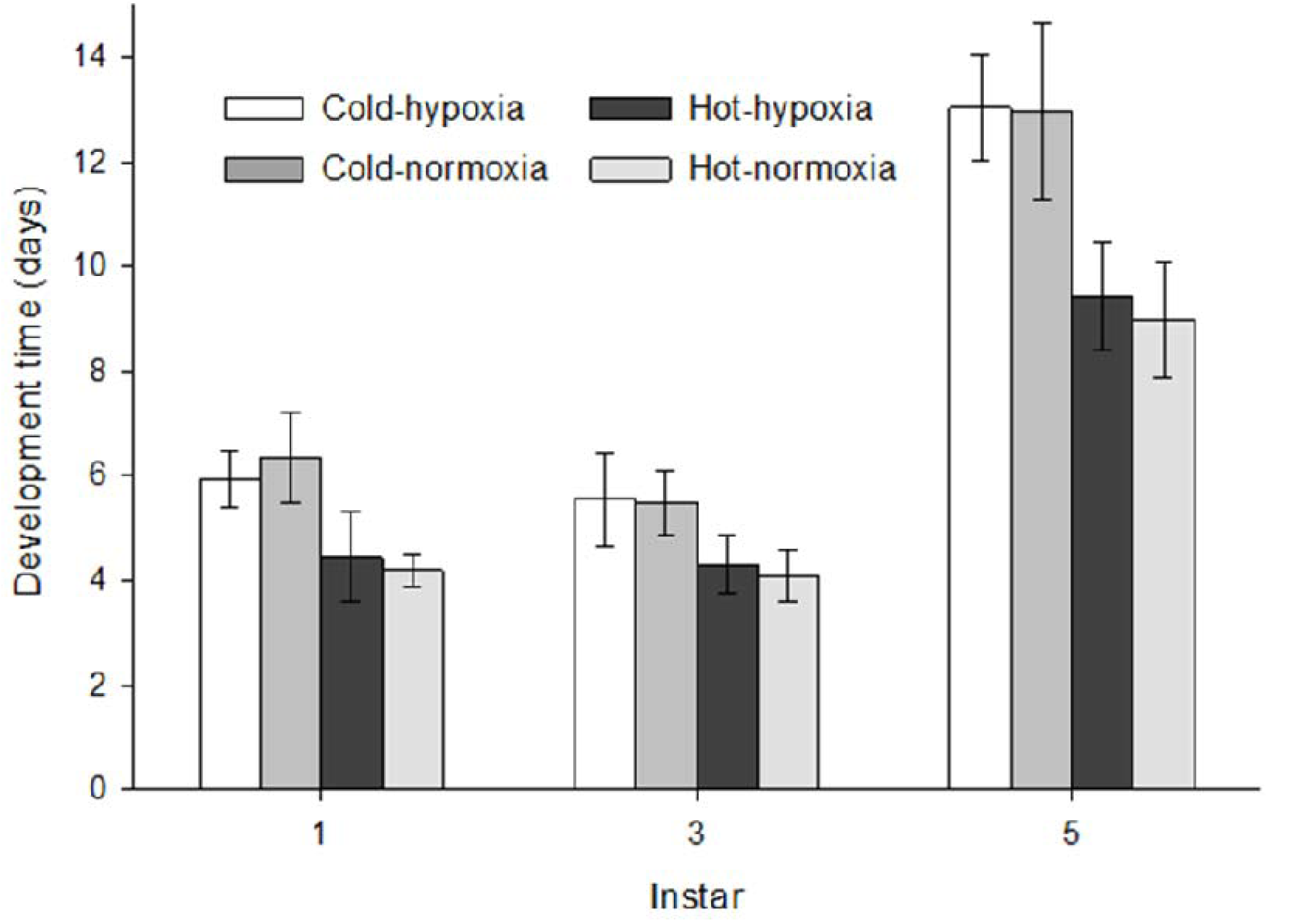
Locusts developed faster at high temperatures. Bars denote the most likely mean of development time, calculated from multimodel averaging, for locusts reared in cold or hot thermal regimes (21-35 or 28-42 °C, respectively) under either hypoxia or normoxia (13 and 21% oxygen, respectively). Error bars represent 95% confidence intervals.

## 4. Discussion

If oxygen supply limits thermal tolerance (Pörtner, 2002; Pörtner, 2010), locusts should have been most susceptible to heat and hypoxia during the early instars because oxygen delivery becomes easier as locusts develop (Greenlee and Harrison, 2004a; Greenlee et al., 2009; Harrison et al., 2005; Hartung et al., 2004; Kirkton et al., 2005; Lease et al., 2006). As predicted, heat and hypoxia interacted to kill more locusts during the first instar than any other instar (Fig. 1). This finding supports the idea that respiratory performance affects an organism’s susceptibility to heat and hypoxia. In a closely related species (*S. americana*), grasshoppers in later instars delivered more oxygen than those in the first instar by ventilating faster and deeper; Greenlee and Harrison (2004a) found that at 35 °C, grasshoppers in the first, third, and fifth instars sustained metabolism down to 13.8, 9.9, and 4.9% oxygen, respectively. Assuming these ontogenetic patterns hold for *S. cancellata*, the hypoxia treatment (13% oxygen) would have limited the metabolism of locusts in the first instar, but this limitation would only have been detrimental at high temperatures. Locusts in the first instars exposed to the cold, hypoxic treatment could have tolerated poor oxygen supply by opening their spiracles for longer or making their tracheolar fluid more permeable to oxygen (Greenlee and Harrison, 1998; Harrison et al., 2006; Wigglesworth, 1983). While this may have been sufficient in cold treatments, we suspect that these compensatory mechanisms were eventually outstripped by the demand for oxygen.

Hypoxia had weak effects on survival and development after the first instar. Locusts in the hypoxia treatment developed at the same rate as those in the normoxia treatment (Fig. 3), contrasting previous observations of *S. americana*, which developed more slowly under 10% oxygen than under normoxia (Harrison et al., 2006). Presumably the level of hypoxia in our experiment (13% oxygen) was not severe enough to impact development. Studies of other insects support the idea that a threshold level of hypoxia reduces performance. For instance, hypoxia influenced the survival and development of mealworm beetles (*T. molitor*) only when oxygen concentration dropped below 15% (Loudon, 1988). Similarly, cockroaches (*Blatella germanica*) reared in an atmosphere of 16% oxygen developed as quickly as those reared in 21% oxygen, while lower oxygen levels increased development time (VandenBrooks et al., 2012). Such thresholds reflect the nonlinear nature of hypoxic stress (Harrison et al., 2017; Rascon and Harrison, 2010), but thresholds may depend on interactions between an organism’s thermal environment and its developmental stage.

In many species of terrestrial insects, heat tolerance remains the same over a large range of atmospheric oxygen concentrations, presumably because terrestrial insects have highly efficient respiratory systems and occupy an oxygen-rich environment (Klok et al., 2004; Lehmann et al., 2019; McCue and De Los Santos, 2013; Shiehzadegan et al., 2017; Youngblood et al., 2019). However, the function of insects can be limited by oxygen supply during metabolically demanding activities, such as flight. For instance, at 35 °C, the flight metabolic rates of American grasshoppers (*S. americana*) and migratory locusts (*Locusta migratoria*) decreased sharply under hypoxia (Rascón and Harrison, 2005; Snelling et al., 2017). If flight performance is limited by oxygen supply, an insect’s ability to fly during heat stress may be limited by its capacity to deliver oxygen to its tissues. This hypothesis is supported by a study on fruit flies (*Drosophila melanogaster*); Teague and colleagues (2017) discovered that a genotype’s ability to fly under hypoxic stress depended on its ability to fly under heat stress, indicating that a common mechanism underlies tolerance to heat and hypoxia. Our findings provide further support for the link between hypoxia tolerance and heat tolerance, more insight into the mechanisms of heat tolerance will come from studying the effects of heat and hypoxia on animals engaged in aerobic activity.

Graham and colleagues (1995) proposed that oxygen delivery is more difficult for larger insects, because diffusion rates decrease as the length of tracheae increase. However, recent evidence suggests that large insects compensate for longer diffusion distances by investing a higher proportion of their body mass into the tracheal system (Kaiser et al., 2007). For example, as grasshoppers (*S. americana*) grow and develop, tracheal volume scales hypermetrically with body mass, increasing the respiratory capacity of older, larger grasshoppers (Greenlee et al., 2009; Hartung et al., 2004; Lease et al., 2006). This functional relationship could explain why oxygen supply limited the heat tolerance of *S. cancellata* in the first instar more severely than it limited the heat tolerance of later instars. In general, oxygen limitation at high temperatures occurs in life stages that primarily rely on diffusion for oxygen delivery (Woods and Hill, 2004). Indeed, oxygen supply limits the heat tolerance of embryonic lizards (Smith et al., 2015), turtles (Liang et al., 2015), and quails (Vimmerstedt et al., 2019). Given that all species must develop a respiratory system at some stage of the life cycle, future work should focus on early stages where the respiratory system lacks the capacity to deliver oxygen seen later in life.

## Acknowledgements

We thank Rick Overson, Arianne Cease, and the entire Cease Lab for providing the locusts used in this study.

